# Flow rapidly replenishes scarce nutrients to promote bacterial growth

**DOI:** 10.1101/2024.12.23.630146

**Authors:** Gilberto C. Padron, Zil Modi, Matthias D. Koch, Joseph E. Sanfilippo

**Affiliations:** Department of Biochemistry, University of Illinois at Urbana-Champaign, Urbana, Il 61801; Department of Biology, Texas A&M University, College Station, TX 77843

## Abstract

In laboratory settings, bacteria grow in static culture with more nutrients than they require. However, bacteria in nature experience flowing environments that are nutrient limited. Using microfluidics and single cell imaging, we discover that flow promotes growth of the human pathogen *Pseudomonas aeruginosa* at surprisingly low nutrient concentrations. In static environments, cells require high nutrient concentrations as they steadily consume non-renewable resources. In flowing environments, cells grow robustly at very low nutrient concentrations that are constantly replenished. Our simulation-guided experiments show how stopping flow halts growth within minutes and varying flow impacts growth gradients across bacterial populations. By precisely delivering nutrients using microfluidics, we learned that cells in flow can grow on glucose concentrations 1,000 times lower than those observed in typical laboratory experiments. The ultralow glucose concentrations sufficient for growth in flow closely align with the affinity of bacterial glucose transporters, suggesting that bacteria have evolved in flowing environments with scarce nutrients. Collectively, our results emphasize the limits of traditional culturing approaches and highlight how depleted environments can unexpectedly support bacterial growth.

## Introduction

Many natural environments are constantly changing due to fluid flow. However, experimental systems typically simplify nature and fail to include flow-driven dynamics. Recent studies using microfluidics have revealed how studying bacteria in flow can shift our understanding of biological processes (1–3). For example, the shear force associated with flow has the counter-intuitive effect on enhancing bacterial adhesion (4–7). In addition, flow physically orients bacteria to move upstream on surfaces (8–10) and flow promotes surface colonization by increasing daughter cell attachment (11, 12). Based on these reports, it now appears essential to include flow in our study of bacterial processes to fully understand how bacteria survive in nature.

Biological flow responses fall into two categories: responses to the shear force associated with flow (2, 4) and responses to flow-driven chemical transport (13, 14). While the impact of shear force on bacterial cells involves stretching or bending (15), flow-driven chemical transport can replenish or wash away small molecules. Logically, the transport of small molecules by flow can amplify or inhibit bacterial processes that depend on local chemical concentrations. For example, flow can amplify the impact of antimicrobials by replenishing molecules faster than cells can remove them (14, 16, 17). In contrast, flow can inhibit quorum sensing by washing away autoinducers faster than they diffuse back to cells (18, 19). In both of these examples, including flow in experiments dramatically changed the results and the interpretation of how bacteria function in natural environments. Thus, using microfluidics to test the impact of flow-driven transport offers a great opportunity to better understand how bacteria interact with their chemical environment.

Bacteria require an adequate supply of nutrients to grow (20, 21). In laboratory experiments, bacteria are typically grown in culture media with an excess of nutrients (22, 23). While complex media allows for robust growth of diverse bacterial species, the use of minimal media allows for precise characterization of nutrient requirements. Minimal media typically consists of a few defined ingredients: water, a carbon source, and various salts which provide elements necessary for growth (24). The presumed advantage of defined minimal media is that all added ingredients are known. By limiting one ingredient at a time, researchers have determined how specific nutrients impact bacterial growth in static culture (25–27). However, it remains unclear how nutrient limitation impacts growth in flowing conditions. Based on the success of studying other chemical transport processes under flow (14, 17, 19, 28, 29), we hypothesize that combining flow and nutrient limitation should yield important new insights.

Using microfluidics, we discover that flow enhances growth of *Pseudomonas aeruginosa* in carbon and nitrogen limited conditions. First, we establish the carbon and nitrogen concentrations required for growth in traditional batch culture. Second, our results reveal that cells in flow can grow at surprisingly low nutrient concentrations. Third, we combined simulations and microfluidic experiments to quantitatively determine how flow modulates spatial and temporal growth gradients. Fourth, we demonstrate that cells in flow can grow with glucose concentrations 1,000 times lower than traditionally observed. Notably, our finding that bacteria can grow on low micromolar glucose levels closely matches the known affinity of bacterial glucose transporters (30, 31), suggesting that bacteria evolved in nutrient depleted flowing environments.

## Results

To begin our study of nutrient limitation in *P. aeruginosa*, we quantified growth in batch culture. We used minimal M9 media with glucose as the sole added carbon source and ammonium chloride as the sole added nitrogen source. To examine how glucose concentration impacts growth, we varied the glucose concentration while keeping all other nutrients constant. From 0.05% to 1% glucose, the growth rate for the early portion of the experiment remained similar (Figure 1A). However, higher glucose concentrations led to higher maximal culture density, while lower concentrations led to lower maximal density (Figure 1A). By varying glucose concentrations over many orders of magnitude, we observed that approximately 0.01% glucose was required to support detectable growth in batch culture (Figure 1B). Similarly, we varied the ammonium chloride concentration while keeping all other nutrients constant. The ammonium chloride concentration impacted the maximal culture density (Figure 1C) and approximately 0.1 mM ammonium chloride was required to support detectable growth in batch culture (Figure 1D). Together, these experiments quantitatively establish the minimum concentrations of glucose and ammonium chloride required to support *P. aeruginosa* growth in batch culture.

**Figure 1.**
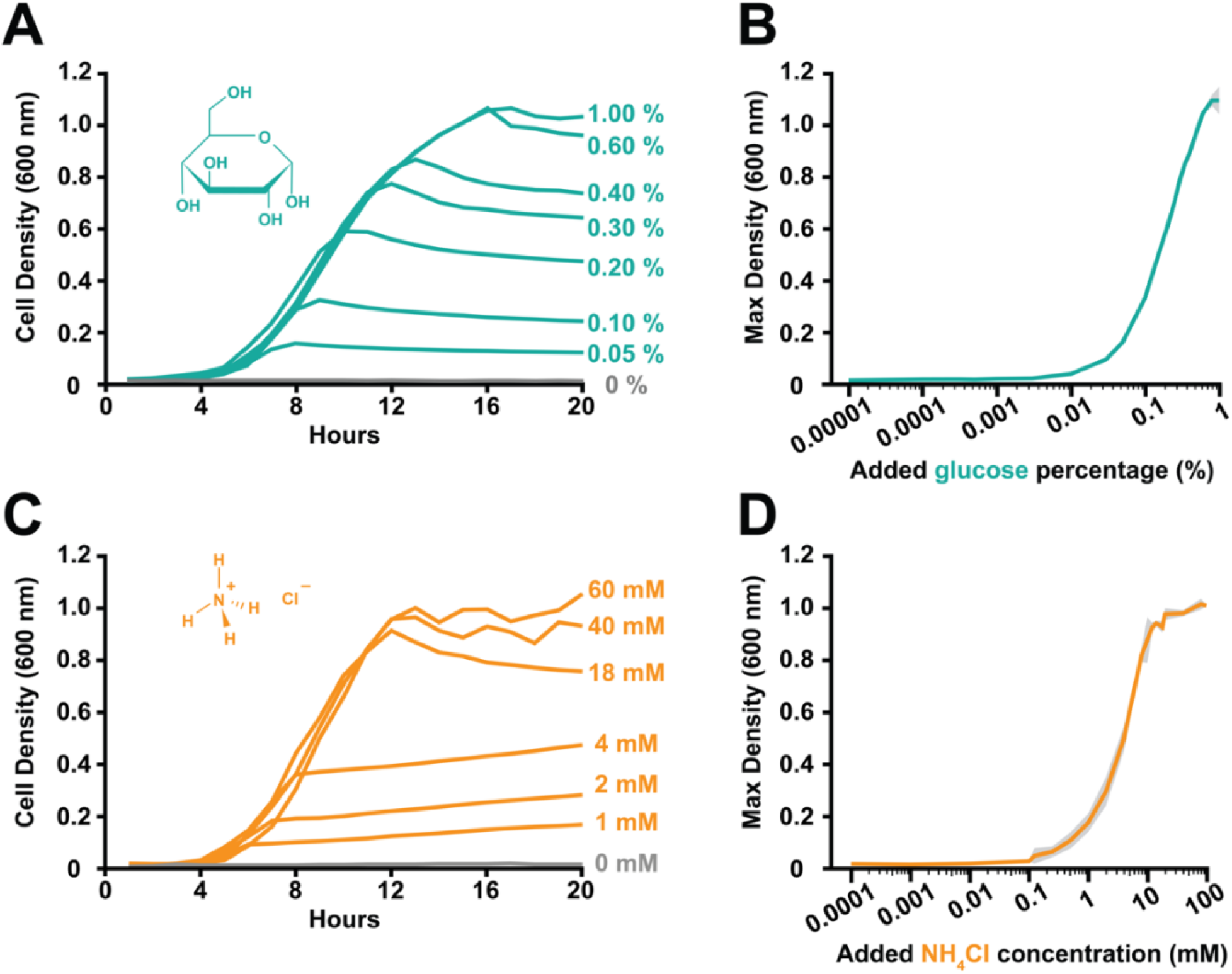
*P. aeruginosa* requires high nutrient concentrations for batch culture growth. **(A)** Growth curves of *P. aeruginosa* cells in M9 minimal media with varying glucose concentrations. Cell density measured by optical density at 600 nm. While increasing glucose impacted maximal culture density, it did not impact growth rate. **(B)** Maximum culture density of *P. aeruginosa* cells grown in varying glucose concentrations. Lines and shading show the average and standard deviation of 3 biological replicates. **(C)** Growth curves of *P. aeruginosa* cells in M9 minimal media with varying ammonium chloride concentrations. Cell density measured by optical density at 600 nm. While increasing ammonium chloride impacted maximal culture density, it did not impact growth rate. **(D)** Maximum culture density of *P. aeruginosa* cells grown in varying ammonium chloride concentrations. Lines and shading show the average and standard deviation of 3 biological replicates.

To examine nutrient limitation with single-cell resolution in flow, we imaged *P. aeruginosa* cells in microfluidic devices. Aiming to determine the minimum glucose concentration required to support *P. aeruginosa* growth in flow, we flowed in M9 media with varying levels of glucose. Surprisingly, our negative control with no added glucose grew robustly (Figure S1). Cells exposed to M9 media with no added glucose grew consistently over an 8 hour period at a shear rate of 800 sec^-1^ (Figure S2). In all our experiments, we represent the flow intensity using shear rate, which describes the flow rate and channel dimensions. Inspired by our observation of growth with no added carbon source, we measured if *P. aeruginosa* cells could grow with no added nitrogen source. Similarly, cells exposed to M9 media with no added ammonium chloride grew consistently over an 8 hour period in 800 sec^-1^ flow (Figure S1, S2).

We interpreted these results to indicate that *P. aeruginosa* cells in flow were capable of growing on carbon and nitrogen contaminants present in our media. Based on these results, we will use the term “carbon limited” to indicate no added carbon source and “nitrogen limited” to indicate no added nitrogen source.

Does flow promote *P. aeruginosa* growth in nutrient limited environments? As growth without added carbon or nitrogen sources was observed in microfluidic devices (Figure S1, S2) but not in batch culture (Figure 1), we hypothesized that flow promotes growth in nutrient limited regimes. By turning flow on and off in microfluidic devices, we tested the flow dependency of growth in carbon or nitrogen limited conditions. In conditions with flow, cells grew in both conditions (Figure 2). However, when flow stopped, cells in both conditions stopped growing (Figure 2). Growth resumed when we restarted flow, indicating that flow promotes *P. aeruginosa* growth in nutrient limited environments. These findings support the hypothesis that flow replenishes nutrients fast enough that cells can continuously utilize them, allowing for growth at lower nutrient concentrations than previously observed.

**Figure 2.**
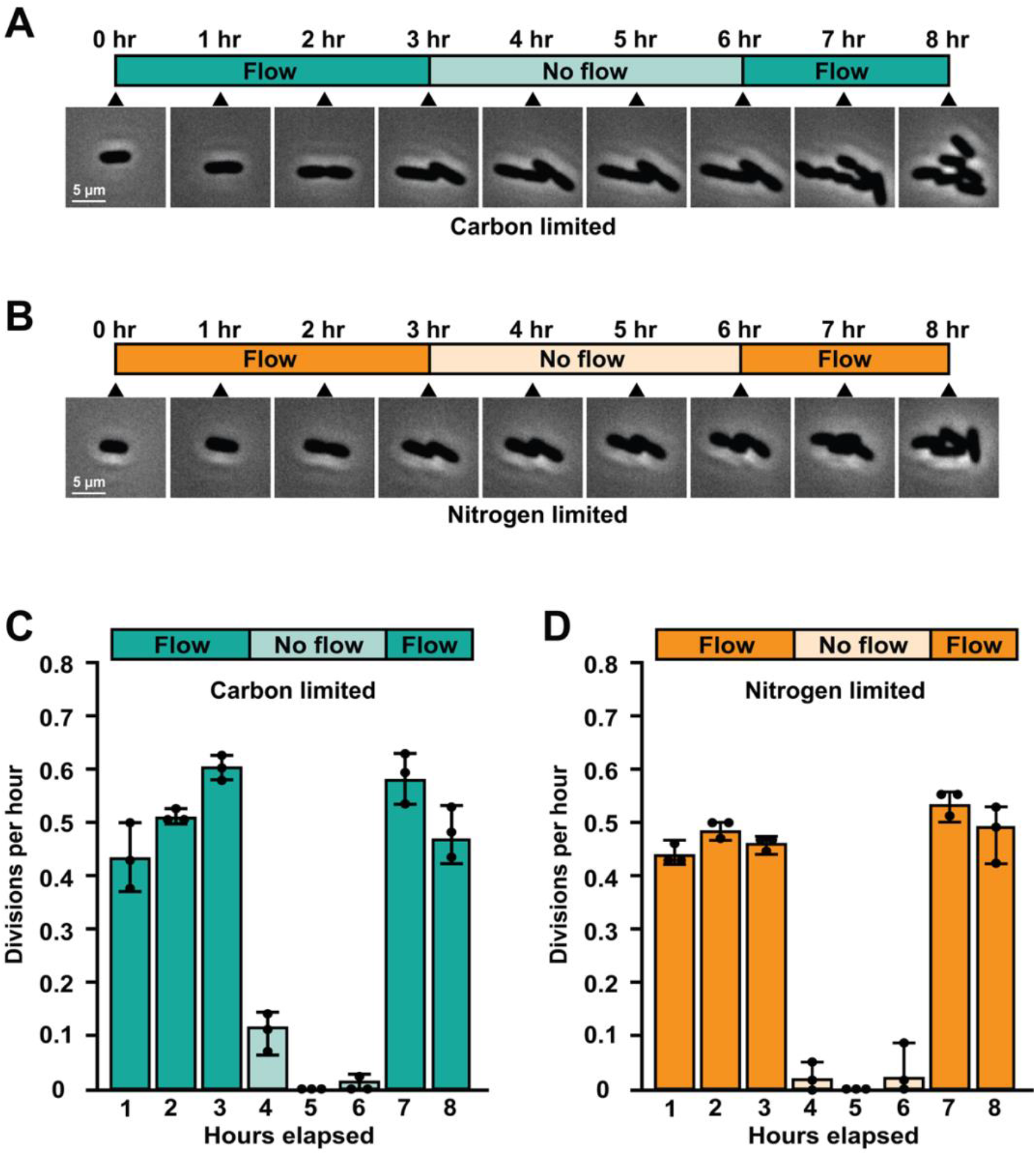
Flow is required for growth with extremely low nutrient concentrations. Timeline and phase contrast images of *P. aeruginosa* during carbon limited **(A)** and nitrogen limited **(B)** growth experiments where flow was turned off and back on. Carbon limited indicates M9 minimal media without an added carbon source and nitrogen limited indicates M9 minimal media without an added nitrogen source. When flow was on, cells could grow in both carbon and nitrogen limited regimes. When flow stopped, cells were quickly unable to grow. Triangles indicate the times at which images were taken. Scale bar on images indicates 5 µm. Quantification of cell division per hour during carbon limited **(C)** and nitrogen limited **(D)** experiments shows the average and standard deviation of 3 biological replicates. For each biological replicate, 30 cells were chosen at random for quantification.

To quantitatively understand the relationship between flow and nutrient limitation, we simulated the flow to no flow transition (Figure 3). Our simulations included three parameters: flow, diffusion, and nutrient removal. By simulating microfluidic channels with flow, we established a baseline nutrient concentration. Then, we stopped flow and quantified how quickly nutrients are depleted from the simulated channel (Figure 3A). For a carbon source the size of glucose, our simulations estimate that approximately 90% of molecules are removed within 30 minutes of stopping flow (Figure 3A). Similarly, for a nitrogen source the size of ammonium chloride, our simulations estimate that approximately 90% of molecules are removed within 15 minutes of stopping flow (Figure 3A). Thus, our simulations generate the testable prediction that available carbon and nitrogen will be depleted within minutes after flow stops.

**Figure 3.**
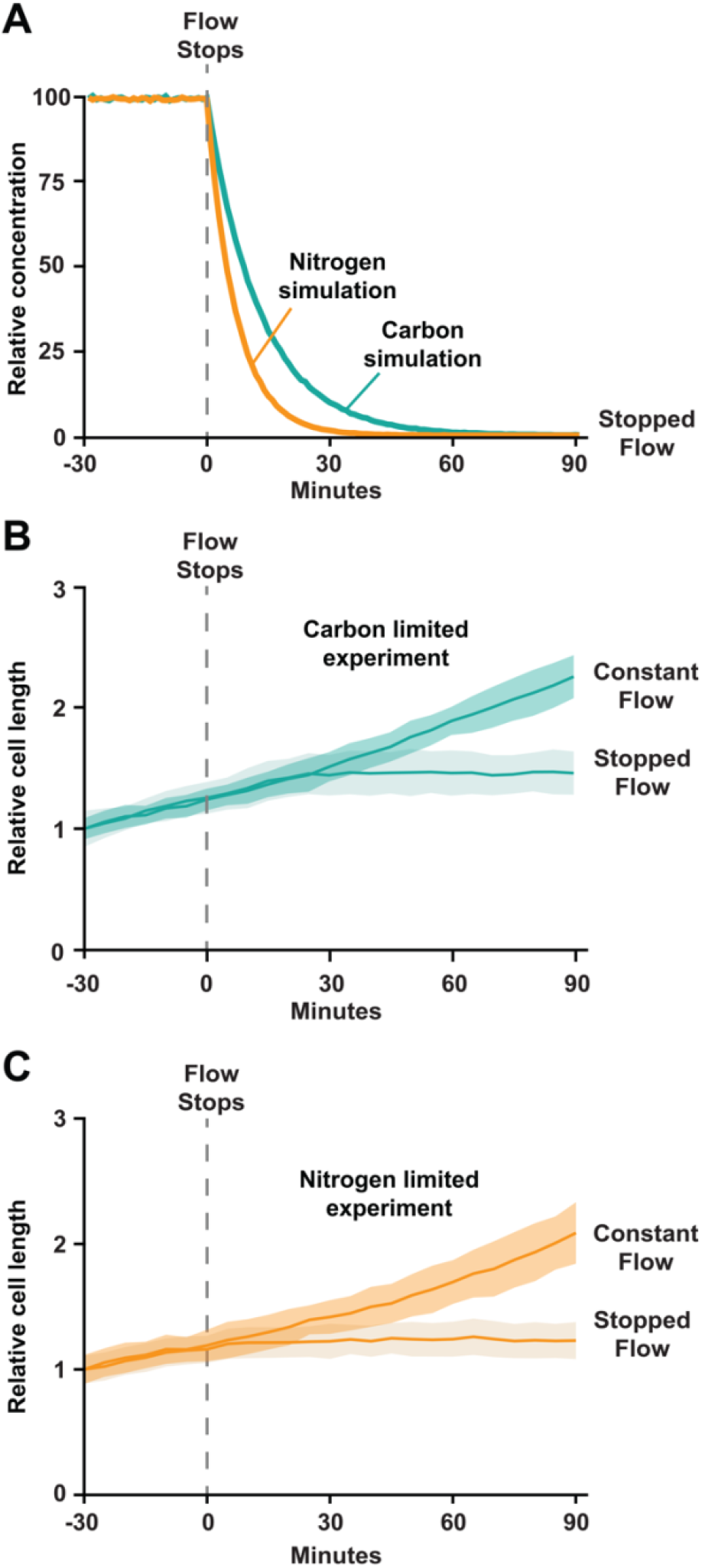
Cells rapidly deplete nutrients and stop growing when flow stops. **(A)** Simulation displaying relative concentration of nutrients before and after flow is shut off. Teal line represents carbon limited simulation and orange line represents nitrogen limited simulation. Quantification of relative cell length (a measure of growth) during carbon limited **(B)** and nitrogen limited **(C)** experiments under constant flow and flow that was stopped. Constant flow indicates flow throughout the experiment, while stopped flow indicates flow that was stopped at the gray dashed line. When flow stops, cells stop growing in less than 30 minutes, which corresponds to the simulated time of nutrient depletion. Lines and shading show the average and standard deviation of three biological replicates. For each biological replicate, 30 cells were chosen at random for quantification.

Guided by our simulations, we experimentally tested the impact of stopping flow on cell growth. To precisely measure growth of cells on short timescales, we quantified the length of cells exposed to constant flow or flow that had just been stopped (Figure 3B, 3C). Cells in carbon limited or nitrogen limited conditions with constant flow elongated at a constant rate (Figure 3B, 3C). In contrast, cells in carbon limited conditions stopped elongating approximately 30 minutes after flow stopped (Figure 3B). Similarly, cells in nitrogen limited conditions stopped elongating approximately 10 minutes after flow stopped (Figure 3C). Our experiments validate our simulations (Figure 3A), demonstrate that cells rapidly stop growing when flow stops, and further support that flow promotes growth by replenishing nutrients fast enough that cells can continuously utilize them.

How does shear rate impact growth in nutrient limited environments? Based on our model that flow promotes growth by rapidly delivering nutrients, we reasoned that shear rate is important. To quantitatively test our reasoning, we simulated a carbon-limited environment in long microfluidic channels with different shear rates. At a low shear rate (80 sec^-1^), short spatial gradients form as molecules are removed before they reach the end of the channel (Figure 4A, 4B). In contrast, at a high shear rate (800 sec^-1^), long spatial gradients form as flow transports molecules deeper into the channel before they can be removed (Figure 4A, 4B). While performing these simulations, we realized that preferential growth at the beginning of the channel may alter the spatial gradient over time. By adding a feedback loop to our simulation where increased growth leads to an increase in the uptake rate (because cells grow and divide), our simulations revealed that spatial gradients become steeper and shift forward over time (Figure S3). Thus, our simulations predict that shear rate has an important role in shaping nutrient gradients across bacterial populations.

**Figure 4.**
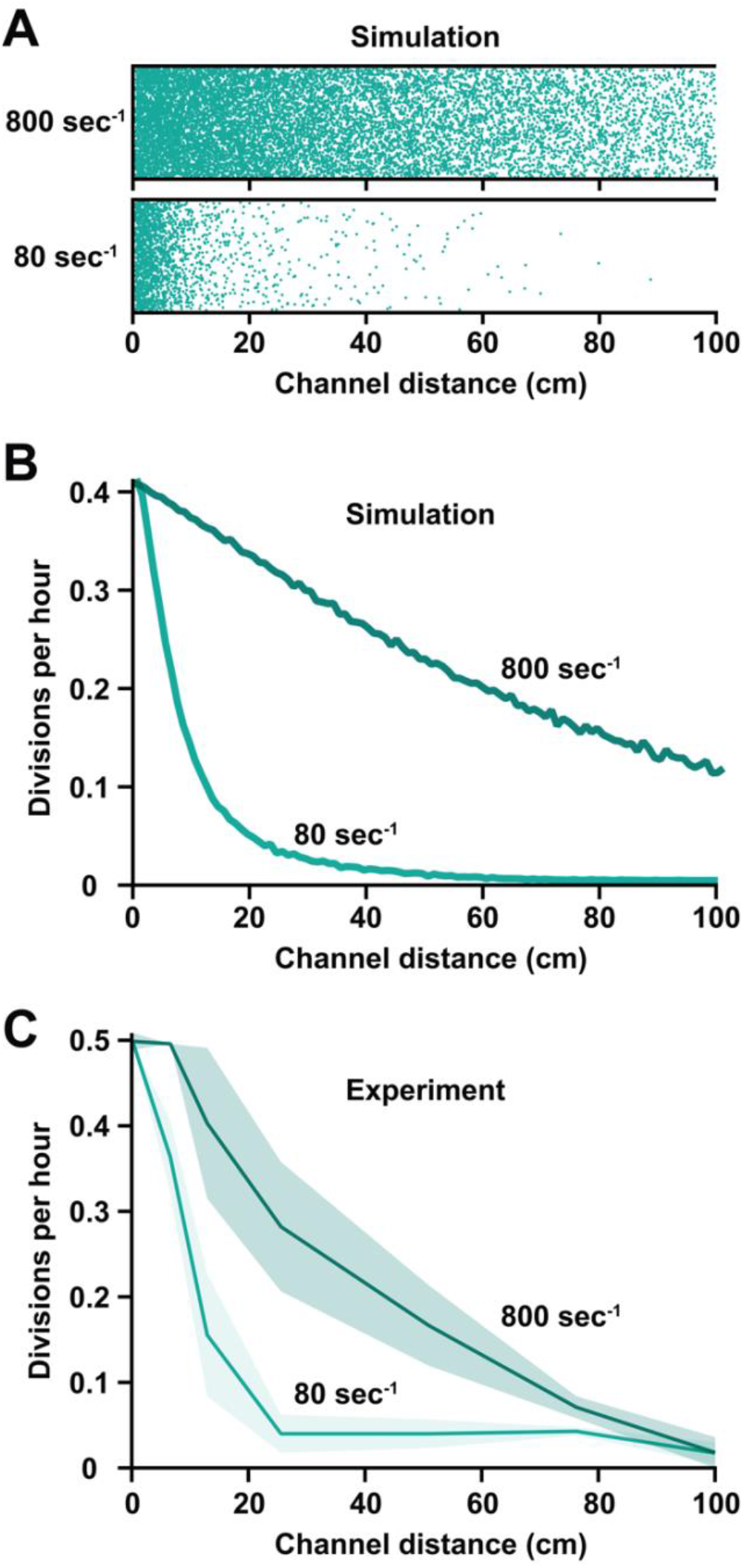
Flow shapes spatial gradients of nutrient availability and growth. **(A)** Visual representation of glucose molecules across simulated 1 meter microfluidic channels at different shear rates. Teal dots represent individual glucose molecules experiencing diffusion in all directions, flow from left to right, and removal when they hit the bottom of the channel. **(B)** Simulated growth gradients based on simulated glucose profiles and the relationship between glucose concentration and growth rate. Faster flow increases the glucose concentration and the growth rate. **(C)** Experimental growth gradients showing the relationship between flow, channel length, and growth rate. Lines and shading show the average and standard deviation of three biological replicates. For each biological replicate, 30 cells were chosen at random for quantification. The experimental results in C closely match the simulated results in B.

To experimentally test the role of shear rate, we measured bacterial growth across long microfluidic channels. Using 1 meter long channels, we quantified growth of *P. aeruginosa* cells in a carbon limited environment exposed to a low shear rate (80 sec^-1^). While cells at the start of the channel grew well, growth decreased quickly as a function of channel length (Figure 4C). At a higher shear rate (800 sec^-1^), we observed that cells grew better and exhibited longer spatial gradients (Figure 4C). The observed spatial gradients for both shear rates were very similar to our simulated spatial gradients (Figure 4B). Analogous shear rate experiments in nitrogen limited conditions revealed similar gradients (Figure S4), indicating that shear rate is generally important during nutrient limitation. Further supporting our simulations (Figure S3), we observed that our spatial gradients become steeper and shift forward over time (Figure S5). Together, our results establish that shear rate has an important role in delivering nutrients into populations, overcoming nutrient limitation, and shaping spatial gradients.

What is the minimum glucose concentration required for growth in flow? As *P. aeruginosa* cells grow in flow with no added carbon source (Figure 2, S1, S2), we hypothesized that cell growth requires very low available carbon. However, as cells were growing on a carbon contaminant, we were unable to quantify how much carbon is required for growth in flow. To overcome this challenge, we flowed our media through microfluidic channels filled with cells at a very high density. At high density, cells depleted the contaminating carbon source and were unable to grow (Figure S5). By collecting and filtering the conditioned media, we precisely determined how much carbon is required for growth in flow (Figure 5). While conditioned media alone could not support *P. aeruginosa* growth in a new channel, conditioned media with 0.00001% glucose added was capable of supporting growth in flow (Figure 5B). In contrast, 0.01% was required to support the same growth in channels without flow (Figure 5C). Thus, our results establish that bacterial cells can grow with 1,000 times lower glucose concentrations in flowing conditions.

**Figure 5.**
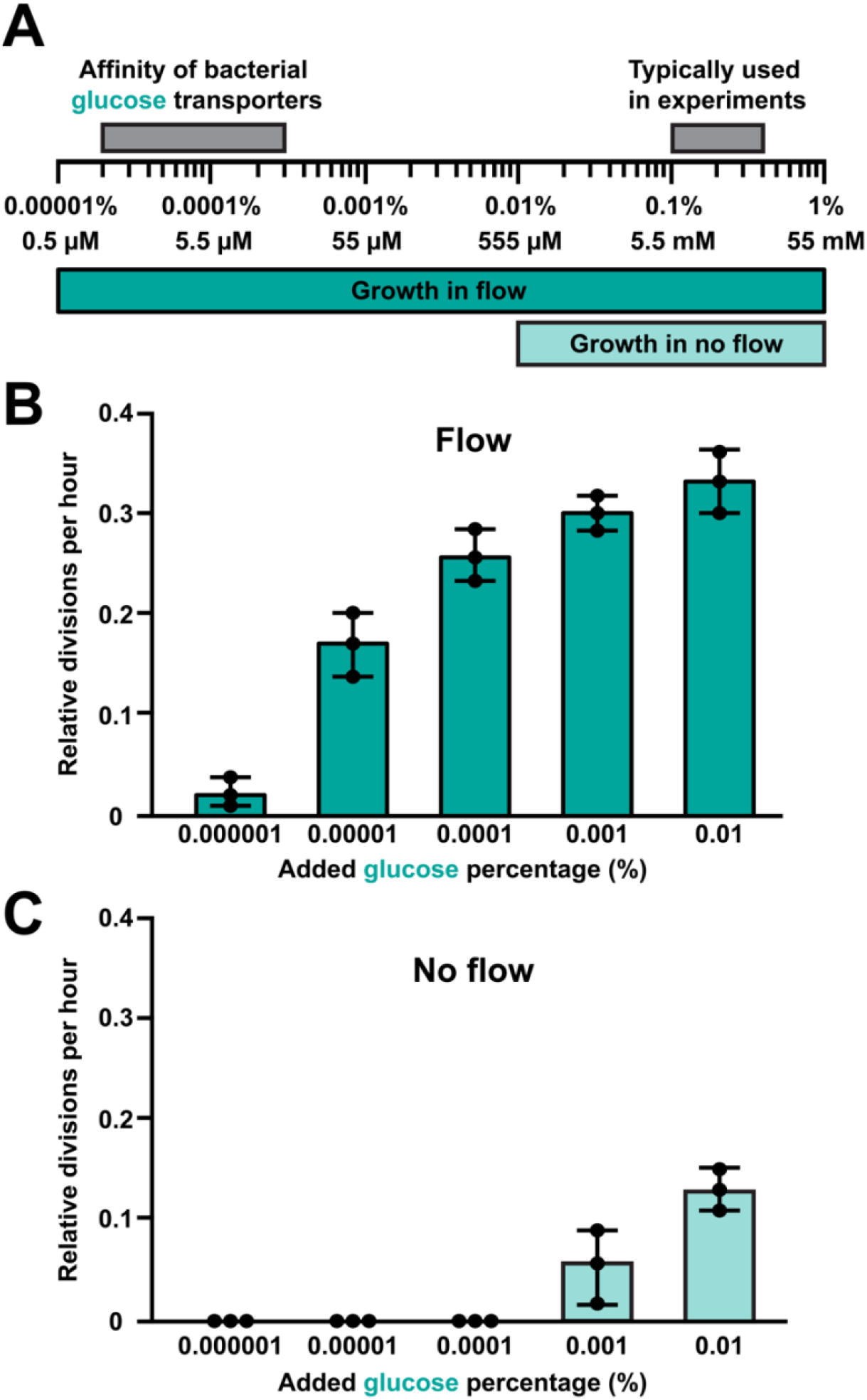
Bacteria grow with 1,000 times lower glucose concentrations in flow. (**A**) Representation of glucose concentrations required for growth in flow and in no flow. For comparison, the glucose concentrations typically used in experiments and the affinity of bacterial glucose transporters was included (30, 31). Cell divisions per hour under flow at a shear rate of 800 sec^-1^ **(B)** or no flow **(C)** using conditioned M9 media with various concentrations of added glucose. M9 media was conditioned by cells to remove contaminant carbon sources. Quantification represents the average and standard deviation of 3 biological replicates. For each biological replicate, 30 cells were chosen at random for quantification.

## Discussion

Here, we report that flow sustains growth of *Pseudomonas aeruginosa* at surprisingly low nutrient concentrations. Using defined minimal media, we demonstrated that cells in batch culture require 0.01% glucose and 0.1 mM ammonium chloride for growth (Figure 1). In contrast, our microfluidic experiments revealed that cells can grow on contaminant levels of carbon and nitrogen when delivered by flow (Figure 2). Combining simulations and experiments, we quantitively determined how quickly cells deplete nutrients in the environment and stop growing (Figure 3). Using long microfluidic channels, we established how shear rate patterns spatial gradients of nutrients and growth across bacterial populations (Figure 4). Finally, we discovered that 0.00001% glucose could support the same amount of growth in flow as 0.01% glucose could support without flow (Figure 5). Together, our results highlight how the interplay between the physical and chemical environment impacts bacterial physiology in nutrient limited regimes.

How do bacteria experience nutrient limitation in nature? While it is difficult to conclusively know how nutrient limitation impacts bacteria in nature, extensive laboratory experimentation using batch culture gives some clues. In batch culture, cells are typically provided with excess nutrients that support growth for many generations. During this phase of exponential growth, cells replicate quickly as they are not limited for nutrients. Eventually, the pool of available nutrients depletes, and the rate of growth slows. At this point, cells enter a nutrient limited regime. However, as this shift occurs rapidly in batch culture, it is technically challenging to quantify the nutrient concentration during the transition to nutrient limitation.

Unfortunately, it is likely that bacteria in nature spend most of their lives transitioning in and out of nutrient limitation (32). For example, bacteria in freshwater (33), marine (34), and host (35) environments are often subjected to nutrient limited flowing environments. Here, we used microfluidics to overcome this challenge and illustrate that bacteria in flow can robustly grow at concentrations that we expected to be limiting. Our unexpected results call for a reevaluation of bacterial nutrient limitation and emphasize that simplified laboratory conditions poorly reflect complex natural environments.

Our discovery that bacteria in flow can grow on extremely low glucose concentrations suggests that bacteria have evolved on the edge of nutrient limitation. To explore this idea further, we compared the glucose concentrations sufficient for growth in flow with the affinity of bacterial glucose transporters. Glucose enters the periplasm of gram-negative bacteria through various non-specific outer membrane porins (30). Periplasmic glucose is then transported into the cytoplasm through specific inner membrane transporters (36–39). In *P. aeruginosa*, the K_m_ for glucose transporters is approximately 8-10 µM (31, 38, 40). In *E. coli*, the K_m_ for glucose transporters is approximately 5-20 µM (30, 41–44). Strikingly, these values closely align with the concentrations sufficient to support growth under flow (Figure 5A). We interpret this alignment to indicate that bacteria have evolved to scavenge low concentrations of nutrients to support growth. Collectively, these results support a model where bacteria in nature can grow by capturing nutrients present at very low concentrations that are replenished by flow.

How does flow-driven chemical transport impact bacteria? Fundamentally, transport is the process of moving something from one place to another. For bacteria in laboratory conditions, nutrient transport is typically driven by diffusion. In closed systems like batch culture, the supply of nutrients by diffusion continues until depletion. However, many natural environments are open systems where transport is facilitated by flow. In these situations, the supply of nutrients by flow is effectively never depleted. Thus, closed diffusion-driven systems require high nutrient concentrations to support many generations of growth, while open flow-driven systems can support many generations of growth on much lower nutrient concentrations. Similar to our findings, recent studies have provided evidence that flow has an important role in many biological processes that depend on small molecules (14, 16, 17, 19, 29, 45). For example, flow’s ability to transport autoinducers suppresses local quorum sensing and facilitates long-distance cell-cell communication (19). Additionally, flow’s ability to deliver antimicrobials faster than cells can neutralize them overcomes bacterial resistance (17). As flow plays a critical role in bacterial growth, cell-cell signaling (19), and antimicrobial resistance (17), it is now clear that experiments should include flow in order to capture the complexities of natural environments.

## Acknowledgements

We thank Anu Sharma, Alex Shuppara, Jessica Palalay, Piyush Sharma, Evan Johnson, Sizhe Chen, Jim Imlay, Wilfred van der Donk, Ben Bratton, and Nick Martin for helpful discussions and comments on the manuscript. This work was supported by grant R35GM155280 to M.D.K. and grants K22AI151263 and R35GM155443 from the National Institutes of Health to J.E.S.

## Contributions

G.C.P., Z.M., M.D.K and J.E.S. designed research. G.C.P., Z.M., and M.D.K. performed research. G.C.P., Z.M., M.D.K. and J.E.S. analyzed data. G.C.P. and J.E.S. wrote the paper.

