## Supplemental Materials for "Flow rapidly replenishes scarce nutrients to promote bacterial growth"

**This includes:**

Materials and Methods
Supplemental Figures S1 to S5

### Materials and Methods

Strains, plasmids and growth conditions

Bacterial strains used in this study are WT PA14 and a Δ*pilA* mutant of WT PA14 (6). All bacterial strains used were plated on LB plates. All *P. aeruginosa* cultures unless explicitly stated were grown at 37°C in M9 media (produced in-house) supplemented with 20% glucose (VWR Life Science) to a final dilution of 0.4% as the carbon source. M9 media was prepared using the Cold Spring Harbor protocol for M9 media preparation freely available from their website. Modifications to the M9 media were made by excluding either glucose or ammonium chloride (VWR Life Science) dependent on experimental condition being tested. Glucose (0-2%) and ammonium chloride (0-200 mM) concentrations were then supplemented into the M9 media as necessary. M9 media was additionally supplemented with 1 M magnesium sulfate (Fisher Chemical), 1 M calcium chloride (Sigma-Aldrich) and adjusted to 7.4 pH using sodium hydroxide for a working solution.

*P. aeruginosa* growth curves

Overnight cultures of *P. aeruginosa* were grown in M9 media as described above. 96-well plates (Avantor) were prepared with addition of glucose (0-2%) and ammonium chloride (0-200 mM). Total media volume was set to 200 µL to ensure a 1:100 dilution of overnight cells (2 µL). Cells were shaken in plate reader (BioTek Synergy S1) at 37°C and optical density measurement were made every hour for 24 hours. Growth curves were derived directly from Gen5 plate reader software using an M9 no cells control as a blank for background subtraction.

Fabrication of microfluidic devices

Microfluidic devices were created and fabricated using soft lithography. Devices were designed on Illustrator (Adobe Creative Suite) and masks were printed by CAD/Art Services. Molds were produced using 100 mm silicon wafers (University Wafer) and were spin coated using SU-8 3050 photoresist (MicroChem). Polydimethylsiloxane (PDMS) chips were plasma treated for bonding onto 60 mm x 35 mm x 0.16 mm superslip micro cover glass (Ted Pella, Inc.). Devices used in concentration and flow to no flow to flow experiments were designed as 7 parallel channels 500 µm wide x 50 µm tall x 2 cm long. Long channel experiments were conducted in devices 500 µm wide x 50 µm tall x 1 meter long. Both devices contained a single inlet and outlet per channel.

Phase contrast microscopy

Timelapse images were captured using a Nikon ECLIPSE Ti2-E inverted microscope using the stock NIS Elements interface. Microscope was equipped with a Nikon 40x Plan Ph2 0.95 NA objective, a Hamamatsu ORCA-Flash4.0 LT3 Digital CMOS Camera, and Lumencor SOLA Light Engine LED source.

*P. aeruginosa* in microfluidic devices

Prior to loading bacterial cultures, outlets were hooked up with tubing for waste outflow (Brain Tree Scientific Polyethylene Tubing; ID 0.015" x OD 0.043"). *P. aeruginosa* was loaded using 10 µL of mid-log (0.4 OD) culture back diluted from overnight cultures 1:100. Experimental media was loaded into 5 mL syringes (BD) and hooked up using standard tubing over a 26-gauge x ½" hypodermic needle (Air-tite Products) to microfluidic device. Syringes were placed on a syringe pump (KD Scientific Legato 210) which was used to vary flow rate (0 – 10 µL/min), which resulted in shear rates from 0 – 800 $sec^{-1}$. For 1 meter microfluidic device experiments, mid-log (0.4 OD) culture was pre-loaded onto channel using syringe pump. 5 mL syringes (BD) were loaded with 1 mL of back-diluted culture and loaded into microfluidic device prior to imaging in a biosafety cabinet (Labconco Logic+). Syringe pump was used to load cells into device at a shear rate of 400 $sec^{-1}$ for 10 mins. 5 minutes of loading was conducted at the inlet of device and 5 minutes at the outlet to most approximately equilibrate the concentration of cells across the device. For some experiments, flow was stopped by turning the syringe pump off. All experiments with 2 cm long devices were imaged 1 cm into the channel. For 1 meter experiments, images were taken at 0.1, 6.25, 12.5, 25, 50, 75, and 97.5 cm distances.

<u>Quantification of *P. aeruginosa* growth</u>

All experiments unless described explicitly followed the same workflow. Timelapse videos were collected every 5 minutes for up to 8 hours. Videos were then analyzed using FIJI (Fiji is just ImageJ) software. 30 cells were picked at random per experimental replicate and tracked for duration of experiment. Divisions were attributed only when cells had fully replicated as "1" and incrementally increased by 1 per replication event (ex. 1 cell divides into 2 (Total: 1 division), 1 cell divides into 4 (Total: 2 divisions) and so forth). Total cell divisions were then averaged for all 30 cells and divided by the length of experiment to obtain divisions per hour. Divisions per hour were averaged across 3 biological replicates and a standard deviation was obtained. All growth experiments were performed with cells lacking *pilA*, which prevented twitching motility and allowed for efficient cell tracking.

<u>Conditioned Media</u>

Conditioned media was prepared by using a 27 cm channel microfluidic device preloaded with a high density (≥1 OD) of *P. aeruginosa* cells. M9 minimal media with no added glucose was then flowed through the 27 cm channel at an 800 $sec^{-1}$ shear rate. The flowthrough of the experiment was collected in a sterile 50 mL tube. After the collection of the flowthrough was completed, the flowthrough was filtered through a 0.22 µm filter Steriflip unit (MilliporeSigma). Conditioned media was then supplemented with glucose as necessary for individual experiments.

<u>Flow simulations</u>

Flow simulations were performed as described previously (12). Previous simulations assumed no change of particle uptake over time. Here, we remove this restriction and explicitly consider cell growth by updating the particle uptake rate based on the particle concentration in the environment during every time loop of the simulation. In brief, we used the experimentally determined growth rate g(c) as a function of glucose concentration c(x) along the channel and the fit function $g(x) = Ac(x)^b/(c(x)^b + c_0^b)$ with the maximum cell division rate of A = 0.535 per hour, the concentration $c_0$ = 0.033 at which g = A/2, and the exponent b = 1.0793. From this, the removal rate r(x, t+Δt) = r(x, t) * (1 + g(x) * Δt) and cell density d(x, t+Δt) = d(x, t) * (1 + g(x) * Δt) along the channel was updated during every simulations step with a time interval Δt.

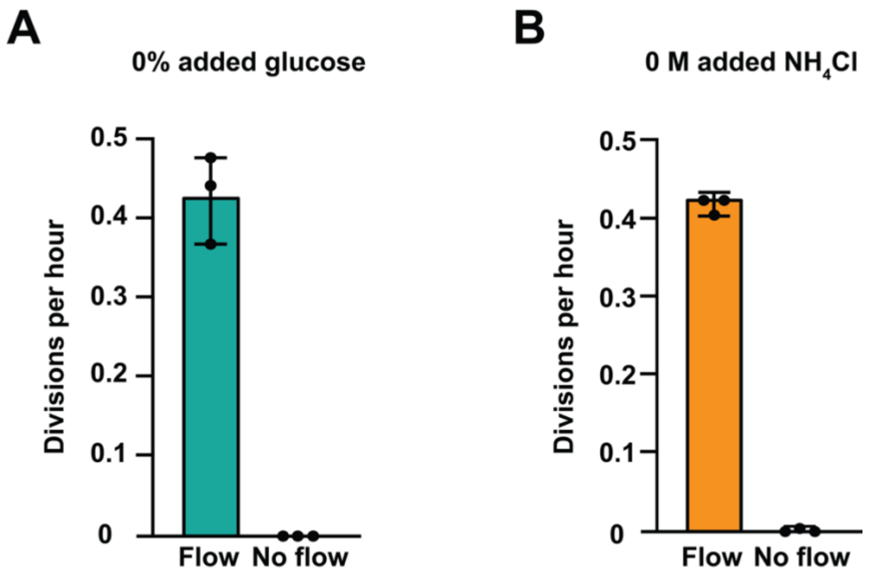


**Figure S1. Contaminant levels of carbon and nitrogen result in flow-dependent growth.** Quantification of *Pseudomonas aeruginosa* growth in flow (800 sec$^{-1}$) and no flow with no added glucose **(A)** or no added ammonium chloride **(B)** in M9 minimal media. Quantification indicates the average and standard deviation of 3 biological replicates. For each biological replicate, 30 cells were chosen at random for quantification.

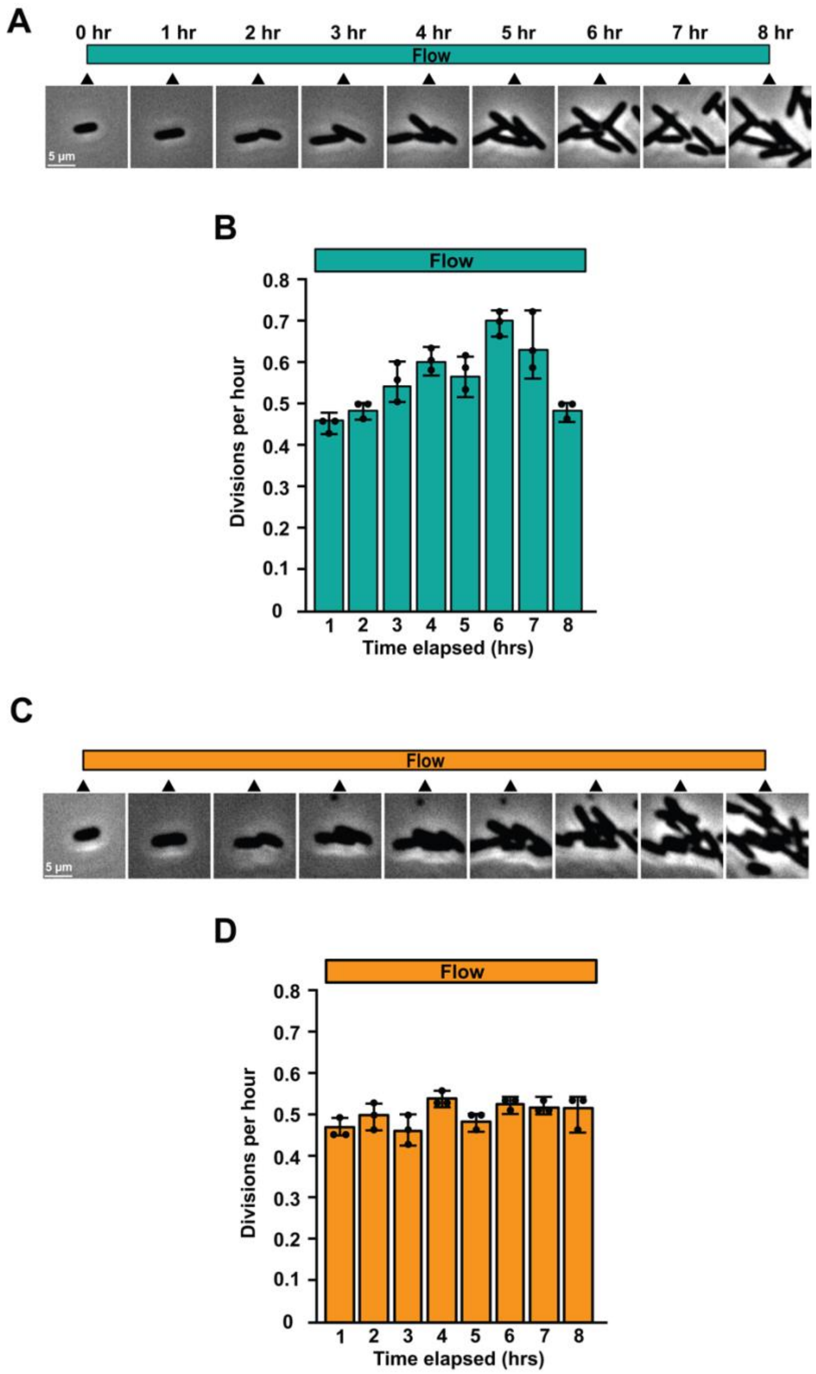


**Figure S2. Flow sustains bacterial growth under nutrient limited conditions.** Timeline and phase contrast images of *P. aeruginosa* during carbon limited **(A)** and nitrogen limited **(C)** growth experiments with flow at a shear rate of 800 $sec^{-1}$. Carbon limited indicates M9 minimal media without an added carbon source and nitrogen limited indicates M9 minimal media without an added nitrogen source. Scale bar on images indicates 5 µm. Quantification of cell division per hour during carbon limited **(B)** and nitrogen limited **(D)** experiments shows the average and standard deviation of 3 biological replicates. For each biological replicate, 30 cells were chosen at random for quantification. Triangles indicate the time at which image was taken.

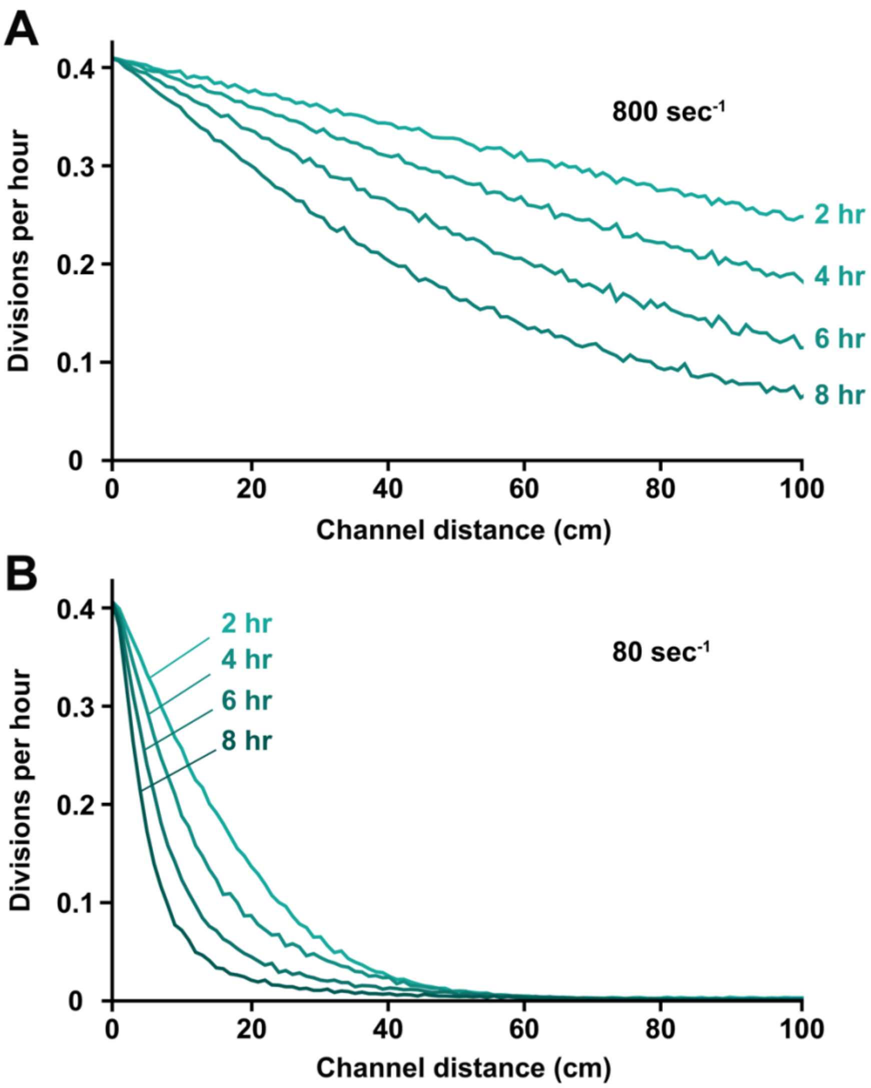


**Figure S3. Simulations predict that spatial gradients shift over time.**
Simulations at 800 sec$^{-1}$ **(A)** and 80 sec$^{-1}$ **(B)** shear rates demonstrating how growth over time impacts spatial growth gradients. As cells grow, their collective ability to remove nutrients increases. In the simulations, a feedback was included where removal increases over time to represent the increase in biomass. Each line represents the rate of growth in divisions per hour at 2 hour time increments. 800 sec$^{-1}$ shear rate generates longer gradients than 80 sec$^{-1}$ because nutrients are delivered more quickly.

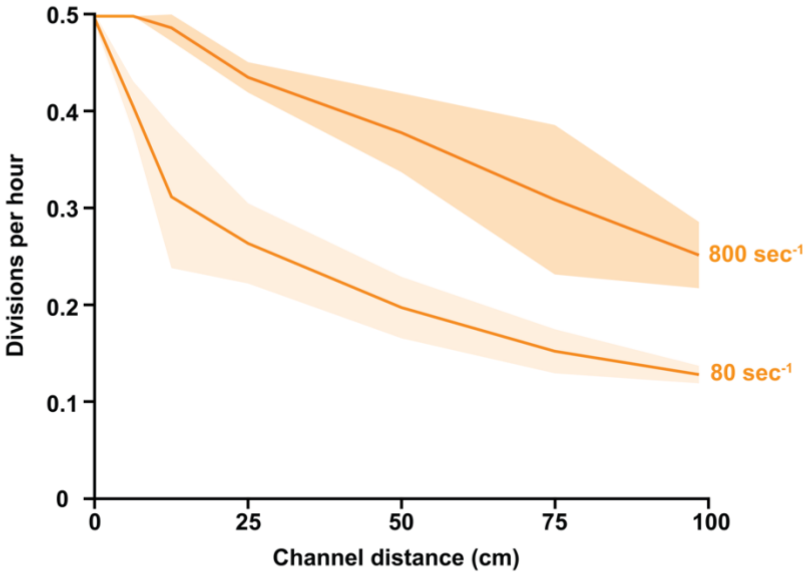


**Figure S4. Flow shapes gradients of nitrogen availability and growth**. Cell divisions per hour under nitrogen limited conditions across a 1 meter channel at different shear rates. For these experiments, M9 media was used with no added nitrogen source. Lines and shading represent the average of three biological replicates and standard deviation. For each biological replicate, 30 cells were chosen at random for quantification.

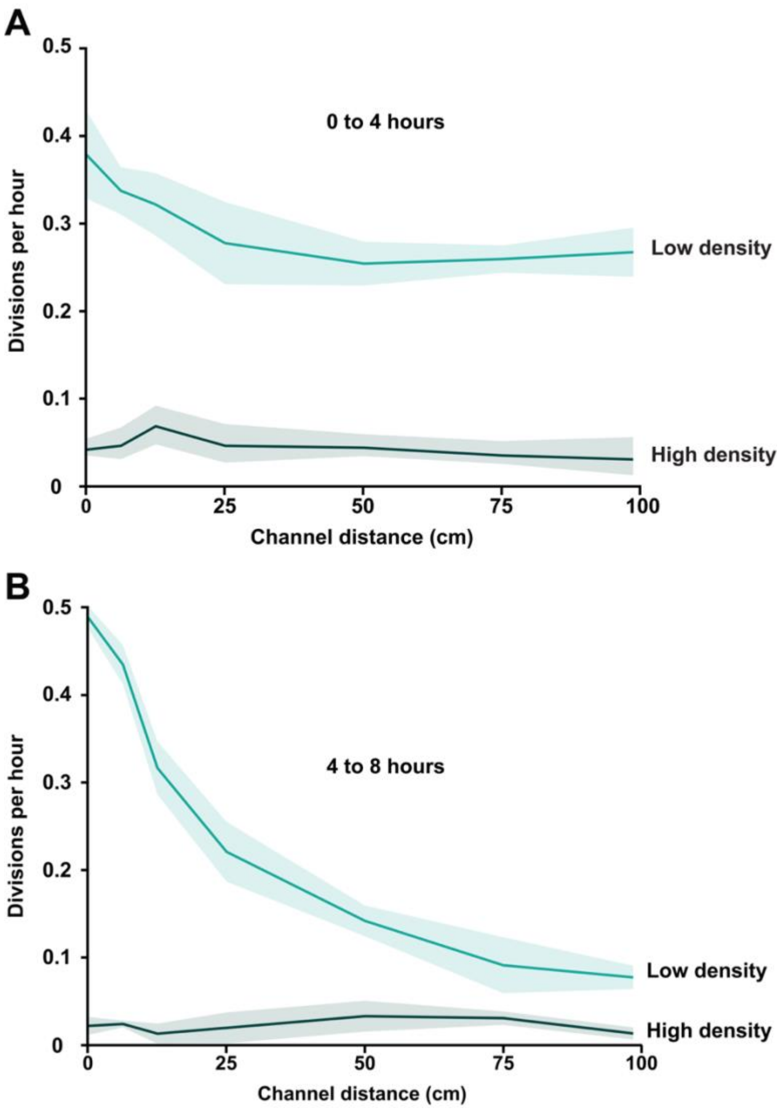


**Figure S5. Cell density impacts growth and shifts growth gradients over time.** Quantification of cell divisions per hour in the first 4 hours **(A)** and second 4 hours **(B)** of an 8 hour carbon limited experiment at both low (10x dilution of a mid-log culture) and high cell density (10x concentrated version of a mid-log culture) at a shear rate of 800 $sec^{-1}$. For these experiments, M9 media was used with no added carbon source. Lines and shading represent the average of three biological replicates and standard deviation. For each biological replicate, 30 cells were chosen at random for quantification.
